# In situ Quantification of Biomolecular Concentration of Cytoplasmic Membraneless Organelles in a Living Cell

**DOI:** 10.1101/2023.05.15.540722

**Authors:** Ren Shibuya, Shinji Kajimoto, Hideyuki Yaginuma, Tetsuro Ariyoshi, Yasushi Okada, Takakazu Nakabayashi

## Abstract

Liquid droplets formed via intracellular Liquid–liquid phase separation (LLPS) are called membraneless organelles and provide enzymatic reaction fields for maintaining cellular homeostasis, while they can be sources of protein aggregates and fibrils, causing neurodegenerative diseases. To understand the nature of intracellular liquid droplets, it is essential to quantify liquid droplets inside a living cell. Here, we performed near-IR fluorescence and Raman imaging to quantify chemical components inside stress granules (SGs) formed via LLPS in living cells under oxidative stress. The Raman images of stressed cells indicate the concentration of nucleic acids in the SGs was 20% higher than surrounding cytoplasm, while the lipid concentration was lower. The intensity of biomolecular C–H bands relative to the water band shows the net concentration of biomolecules was almost the same inside and outside the SGs, indicating intracellular droplets are not highly condensed, but the crowding environments are similar to the surroundings.

**TOC GRAPHICS:** 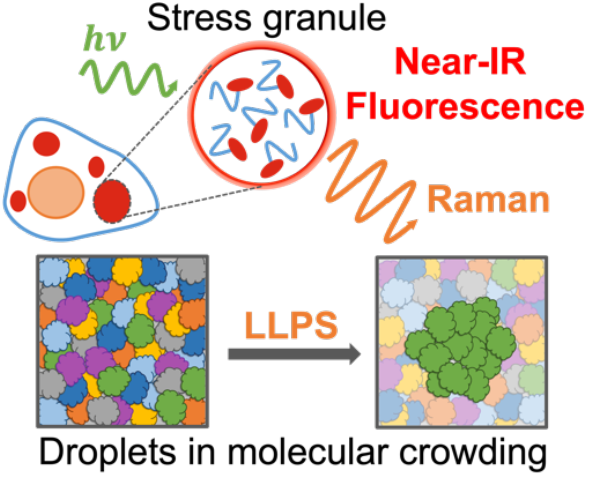

## Introduction

Liquid–liquid phase separation (LLPS), in which a multi-component solution separates into dense and sparse liquid phases, plays a critical role in the compartmentalization of cells.^1-5^ Intracellular liquid droplets formed via LLPS are sometimes called membraneless organelles and provide a unique and transient reaction field for intracellular events.^6,7^ Nucleoli, the factory for ribosome production inside the nucleus, Cajal bodies for RNA retention, and P-bodies for mRNA turnover, are examples of membraneless organelles. On the other hand, liquid droplets have been suggested to be the precursors of the aggregation and fibril formation of proteins, resulting in the pathogenesis of neurodegenerative diseases.^8-10^ Understanding the properties of liquid droplets formed via LLPS is thus crucial for comprehending the various events in cells and the pathogenesis of neurodegenerative diseases.

Quantitative understanding of the chemical components and their concentrations in a droplet is important for elucidating the properties and functions of liquid droplets. Fluorescence imaging is a powerful tool with high spatial resolution and sensitivity to identify liquid droplets, however, it only reveals the presence of labeled molecules, not whether unlabeled molecules are present in the droplet. Photobleaching of fluorescence tags also complicates the concentration quantification of the target molecule. Intracellular droplets can be considered one of the molecular crowding environments representing highly crowded states with various biomolecules in cells; however, it is difficult to discuss the differences in the crowding environment inside and outside droplets, since there is no direct method to quantify liquid droplets and the surrounding intracellular environment in a living cell.

Raman imaging has attracted much attention as a promising method for visualizing the chemical components inside a living cell in a label-free and non-invasive manner.^11-13^ Raman imaging using different Raman bands offers the simultaneous visualization of the distribution of various molecules. Furthermore, the concentration of molecules can be evaluated quantitatively using a water Raman band as an internal standard.^14,15^ This technique allows the concentration determination of biomolecules in a single droplet in buffer solutions and can also be applied to quantify local crowding environments in a living cell.^16,17^

The application of Raman measurements to droplets in buffer solutions has been reported in recent years;^14,15,18-21^ however, its application to actual intracellular droplets is very limited. The drawback of Raman measurement is the low scattering cross section resulting in the low signal level. The large number of small Raman bands derived from various biomolecules in cells complicates the analysis and makes imaging of small cellular organelles difficult. Therefore, except for nucleoli,^22^ observation of intracellular liquid droplets with Raman imaging has been conducted only on fixed cells.^23^

In this study, we combined Raman and near-IR fluorescence imaging to observe and quantify stress granules (SGs) formed in living cells upon oxidative stress (Figure S1 in the Supporting Information). SGs are transient liquid droplets formed as a result of cellular response against various stresses.^24,25^ SGs contain mRNA and various proteins, and are regarded as membraneless organelles, having an essential role in mRNA metabolism and translational control under stress.^26,27^ Besides, SGs have also been implicated in various diseases.^28-30^ Despite the widespread research, quantitative analysis of intracellular SGs has not been explored due to the lack of non-invasive assays. To observe SGs on a Raman microscope, we utilized cells transfected with iRFP-G3BP1, where G3BP1, one of the scaffold proteins of SGs, is labeled with the near-IR fluorescent protein iRFP713. iRFP labeling offers the determination of the position of SGs in living cells without interfering with Raman imaging with visible laser excitation (Figure S2). In this study, we obtained near-IR fluorescence images and determined the position of SGs in living cells, and then measured Raman imaging of SGs to perform in situ quantification of the components in a single SG and the surrounding cytoplasmic regions. We compared the crowding environments in SGs and cytoplasm and discussed the nature of intracellular droplets in terms of the crowding environment.

## Results and Discussion

### Concentration of Biomolecules in Stress Granules

Near-IR fluorescence and the corresponding Raman images of HEK293 cells with and without oxidative stress were shown in Figure 1 (another example is shown in Figure S3). In the control cells, the near-IR fluorescence showed almost uniform intensity throughout the cytoplasmic region, indicating that G3BP1 was distributed homogeneously in the cytoplasm (Figure 1A). On the other hand, in the near-IR fluorescence images of the oxidative-stressed cells, bright spots were observed in the cytoplasm, which can be attributed to SGs (Figure 1E and Figure S3A).

**Figure 1.**
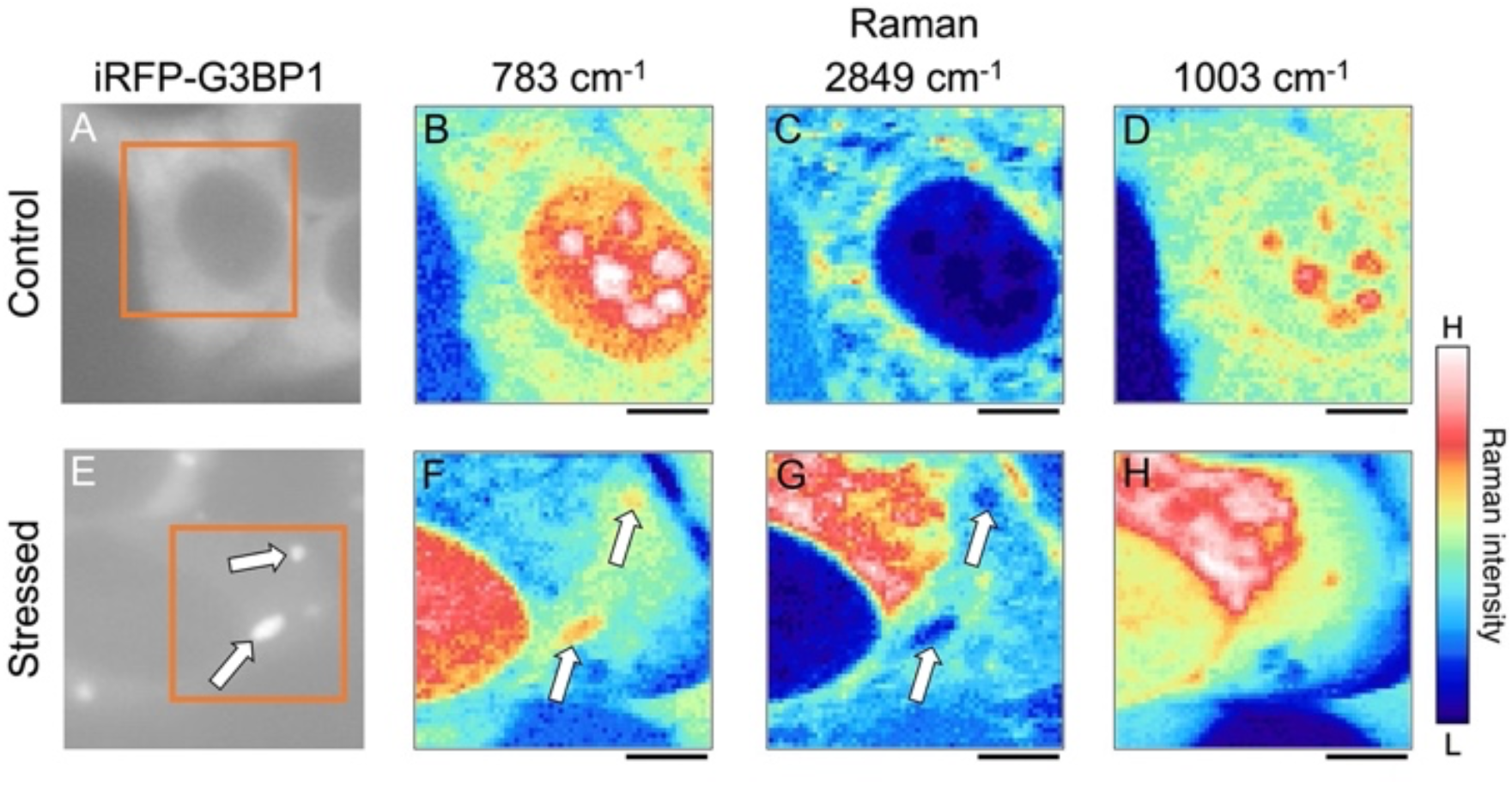
Near-IR fluorescence (A, E) and the corresponding Raman images of control (B-D) and stressed (F-H) cells. Each Raman image was obtained by plotting the Raman intensity of the pyrimidine ring band (783 cm^-1^) (B, F), the CH2 symmetric stretching band (2849 cm^-1^) (C, G), and the phenyl ring breathing band (1003 cm^-1^) (D, H). Orange boxes in the fluorescence images show the region of Raman imaging. White arrows in the images indicate stress granules. Scale bar: 5 μm.

Raman images were obtained by mapping the integrated intensities of Raman bands at 783 cm^-1^ (pyrimidine ring), 1003 cm^-1^ (phenyl ring of phenylalanine), and 2849 cm^-1^ (CH2 symmetric stretching of lipids). A broad baseline was subtracted from each Raman spectrum to obtain Raman images. The Raman images of the stressed cell at 783 and 2849 cm^-1^ exhibited structures resembling SGs in the near-IR fluorescence image (as indicated by arrows in Figure 1E, F, G and Figure S3A, B, C). The Raman intensity at 783 cm^-1^ was higher and that at 2849 cm^-1^ was lower in comparison to the surrounding area, indicating that the concentration of nucleic acids in the structured areas was higher than in the surrounding cytoplasmic region, while the concentration of lipids was lower. The obtained result was consistent with the expected components of SGs, and we conclude that Raman images of the SGs in living cells have been successfully obtained by combining near-IR fluorescence and Raman measurements. In contrast, the Raman image of the phenylalanine band at 1003 cm^-1^ did not show structures assignable to the SGs (Figure 1H and Figure S3D), suggesting that the concentration of proteins in the SGs was comparable to the surrounding area.

We extracted Raman spectra of cytoplasm, nucleoplasm, nucleoli, and SGs from Raman images of stressed and control cells. Figure 2 shows the averaged Raman spectra of the SGs and the surrounding cytoplasmic region in the stressed cells and their difference spectrum (SGs – cytoplasm). The averaged Raman spectra were normalized with the integrated intensity of the water O–H stretching band (3200 -3570 cm^-1^) (Figure 2A), allowing us to discuss the concentration of biomolecules relative to the water concentration. In the fingerprint region of the difference spectrum (Figure 2B), negative peaks were observed at 1737, 1656, 1433, and 715 cm^-1^, assigned to lipids, and positive peaks were at 1484, 1336, and 783 cm^−1^, assigned to nucleic acids (Table S1). These negative and positive peaks confirm that the interior of the SGs is richer in nucleic acids and poorer in lipids than the surrounding cytoplasm.

**Figure 2.**
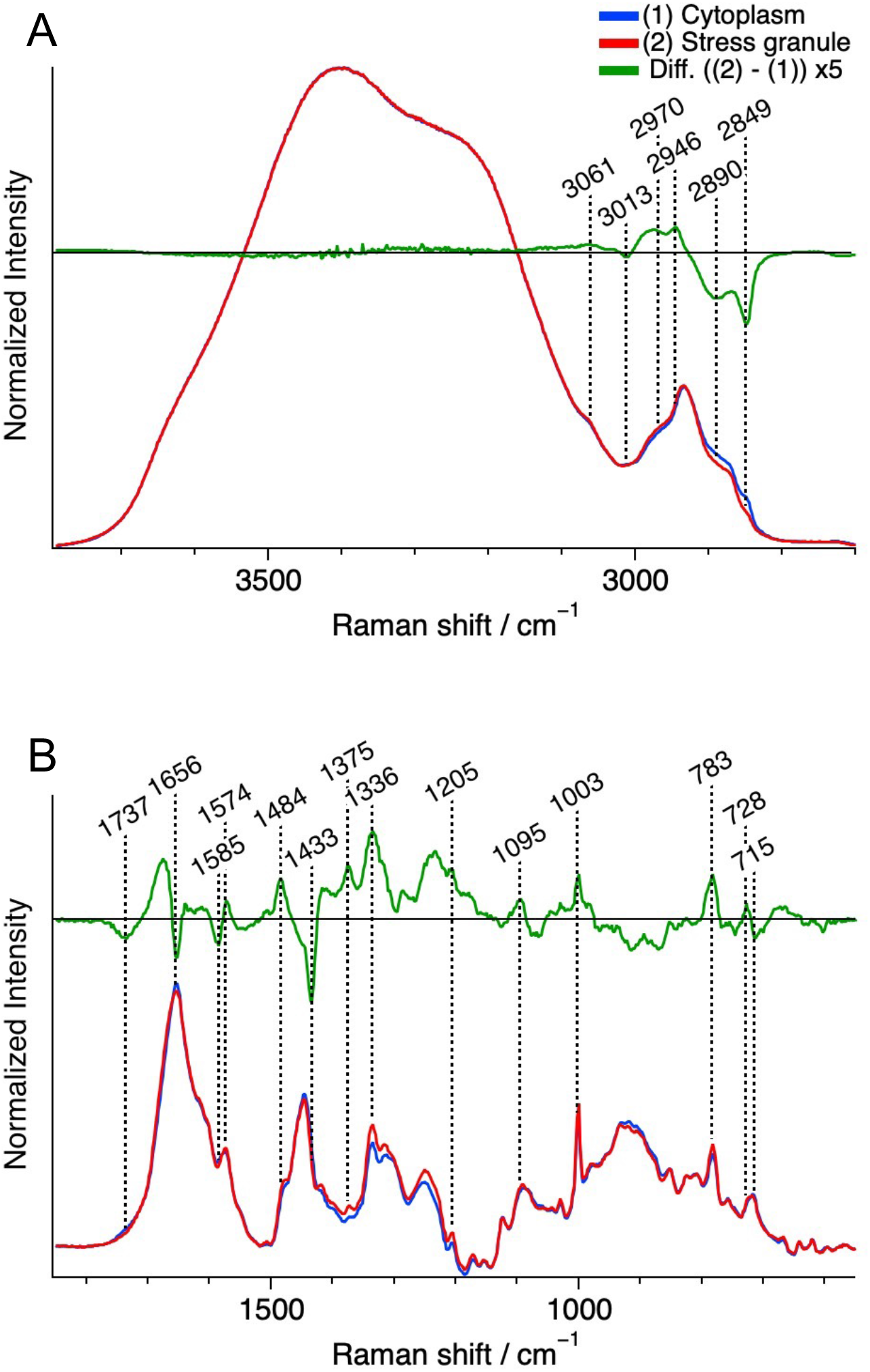
Average Raman spectra of cytoplasm (blue, *n* = 16), stress granules (red, *n* = 12) in stressed cells, and their difference spectrum (granule – cytoplasm) (green) in the C–H and O–H stretching band region (A) and the fingerprint region (B)

A positive peak appeared at 1003 cm^-1^ in the difference spectrum, indicating that the concentration of proteins in the SGs is higher than that in the surroundings. A pair of positive and negative peaks were found in the higher and lower wavenumber regions, respectively, around the protein amide I band (1650 -1700 cm^-1^). This differential-shaped peak can be ascribed to the higher concentration of proteins and the lower concentration of lipids having the C=C stretching band at 1656 cm^-1^ in the SGs. The positive peak in the amide I region in the difference spectrum also indicates that the protein concentration in the SGs is higher than in the surroundings. However, both the positive peak intensities at 1003 cm^-1^ and in the amide I region in the difference spectrum were only about 5% of the corresponding intensities in the original spectra. This result indicates that the difference in the total protein concentration was small between the SGs and the cytoplasm. In the C–H stretching region (2800 - 3000 cm^-1^), the intensity in the low and high wavenumber sides increased and decreased, respectively, in the difference spectrum (Figure 2A). The C–H stretching band of lipid molecules is known to have larger contribution on the low wavenumber side, which is consistent with the result in the fingerprint region that the amount of lipids decreased in the SGs. To summarize, Raman imaging analysis has shown in situ that RNA and certain proteins are assembled and lipids are eliminated during the formation of the SGs.

The intensity of the C–H stretching bands reflects the concentration of biomolecules in the cell. As shown in Figure 2A, the difference in the integrated intensity of the C–H stretching bands between the SGs and the cytoplasm was small, less than 5% (Figure S4). This result means that the total concentration of biomolecules was similar between inside and outside the SGs; namely, the interior of the SGs formed via intracellular LLPS after oxidative stress was not highly condensed, and the crowding environment inside the SGs was nearly the same to the outside. If there is a difference in the concentration of biomolecules, the difference should be reflected in the water density inside and outside the SGs. As determined from the intensity of the O–H stretching band of the intracellular region relative to that of the surrounding medium (Figure S5), there was no significant difference in the water density in the SGs and the cytoplasm. These results indicate that the difference in the total biomolecular concentration was small between the SGs and the cytoplasm.

Results from in vitro experiments on LLPS show that the inside of liquid droplets in buffer solution is highly concentrated with specific biomolecules.^14,15^ This result seems to differ from the present result showing a small difference in biomolecule concentration between the SGs and the cytoplasm in living cells. The difference between in vitro and in vivo comes from the difference in the environment surrounding droplets. The intracellular environments are highly crowded with a variety of biomolecules.^31,32^ Certain molecules, like G3BP1, will be concentrated nearly 1000-fold in the SGs, similar to liquid droplets in buffer solution; however, at the same time, other molecules will be excluded from the droplets, so that the net biomolecular concentration (i.e., the crowding environment) will be almost the same inside and outside the SGs (Figure S6). Note that the present result is consistent with the result of refractive index measurements of SGs,^33^ in which the refractive index inside SGs was similar to that of the outside.

### Stress-Induced Changes in Concentration of Biomolecules

We next compared the Raman spectra of cytoplasmic regions with and without the oxidative stress to investigate the stress-induced change in the concentration of intracellular biomolecules (Figure 3). A negative broad band in the difference spectrum (stressed – control) was observed in the C– H stretching band region (Figure 3A), indicating that the net concentration of biomolecules decreased with the oxidative stress. In the difference spectrum in the fingerprint region (Figure 3B), negative peaks were shown at 1737, 1656, 1445, and 715 cm^-1^, which are attributed to lipids, and 1656, 1336, and 1003 cm^-1^, which are attributed to proteins. These negative peaks indicate that both lipids and proteins in the cytoplasm decreased with the oxidative stress. The magnitude of the decrease was about 10%. The shape of the difference spectrum differs from the original spectrum, indicating that the concentration decrease was not simply due to the increase in the cell volume.^34^ The obtained concentration decrease can be ascribed to the cellular responses to the oxidative stress, such as the decomposition of lipids under the stress and the degradation of oxidized proteins by autophagy. The oxidation and degradation of liposome proteins will also reduce the amount of newly formed proteins, decreasing total protein concentration.

**Figure 3.**
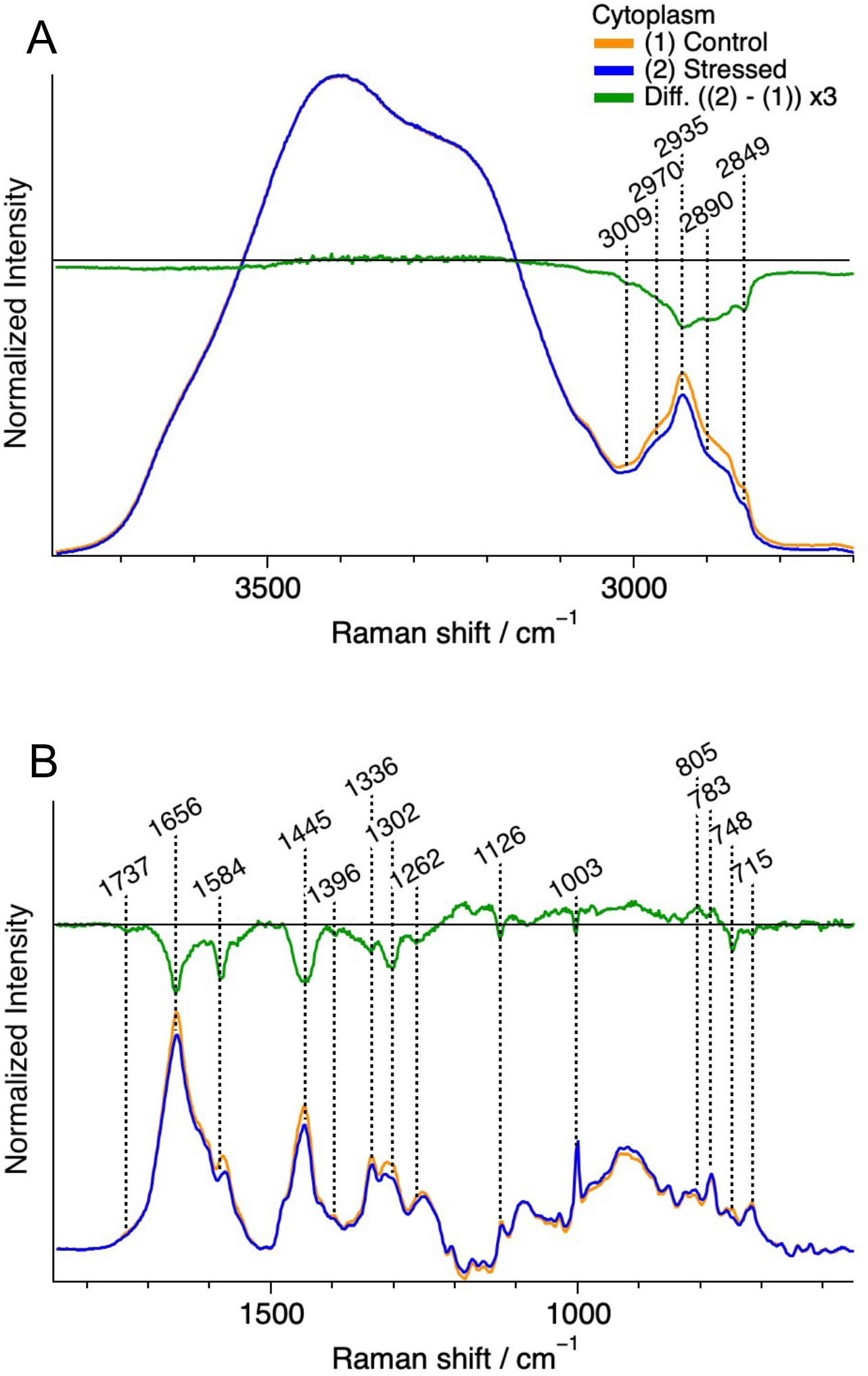
Average Raman spectra of cytoplasmic regions in control (yellow, *n*=12) and stressed cells (blue, *n*=16), and their difference spectrum (stressed – control) (green). Each spectrum was normalized with the integrated intensity of the O–H stretching band (3200 − 3570 cm^-1^). For the fingerprint region (B), a baseline was obtained by fitting with a fifth-order polynomial and subtracted from each Raman spectrum.

Negative peaks were observed at 1584, 1126, and 748 cm^-1^, all of which are assigned to the resonance Raman bands of the reduced form of cytochrome *c*. The decrease in the reduced cytochrome c may come from stress-induced mitochondrial dysfunction. Mitochondria play an important role in lipid homeostasis, and mitochondrial dysfunction may also contribute to the decrease in the lipid concentration. The changes in the intensity of the C–H stretching band region in the nucleoplasm (Figure S7) and nucleoli (Figure S8) were less than a third of the change in the cytoplasm, implying that the environment inside the nucleus was rather maintained against the oxidative stress.

### Quantification of Concentration of Nucleic Acids

We performed the in situ evaluation of the concentration of nucleic acids in the SGs and other regions. The pyrimidine band at 783 cm^-1^ was used to obtain the concentration of nucleic acids. The O–H stretching band of water provides an intensity standard. We first obtained Raman spectra of RNA aqueous solutions (Figure S9), and plotted the intensity of the pyrimidine band relative to that of the O–H stretching band (3200 - 3570 cm^-1^) as a function of the RNA concentration, providing the calibration line (Figure S10). We then calculated the intensity ratio of the pyrimidine band in the each cellular compartment to the O–H stretching band of extracellular water and converted the intensity ratio to the concentrations of nucleic acids using the calibration line. Note that the water density in a cell is not uniform and the intensity of the O–H stretching band of a cell slightly differs from that of buffer solution, making it difficult to use the Raman band of intracellular water as an intensity standard (Figure S5).

Figure 4 shows the plots of the nucleic acid concentration evaluated for different cellular regions in living cells with and without the oxidative stress. The average values are summarized in Table 1. The concentrations of nucleic acids in the cytoplasm, nucleoplasm, and nucleoli in control living cells were evaluated to be around 14, 18, and 28 mg/mL, respectively. These values are comparable with the previous report of stained cells using deep-UV absorption microscopy.^35,36^ The concentration changes of nucleic acids were small after the oxidative stress in the cytoplasm, nucleoplasm, and nucleoli. The concentration of nucleic acids in the SGs was evaluated to be 19 mg/mL, which is 20% higher than the surrounding cytoplasmic region and similar to that in the nucleoplasm. The present result quantitatively shows that nucleic acids were localized to some extent in the SGs, while the concentration of nucleic acids was almost maintained in other regions after the oxidative stress.

**Table 1.** Average nucleic acid concentration in each cellular region of control and stressed cells. Outliers in Figure 4 are excluded from the calculations.

| Cellular region | Nucleic acids concentration / mg mL<sup>-1</sup> |  |
| --- | --- | --- |
|  | Control | Stressed |
| Cytoplasm | 14.1 | 15.4 |
| Nucleoplasm | 18.4 | 19.4 |
| Nucleolus | 28.2 | 27.6 |
| Stress granule |  | 18.8 |

**Figure 4.**
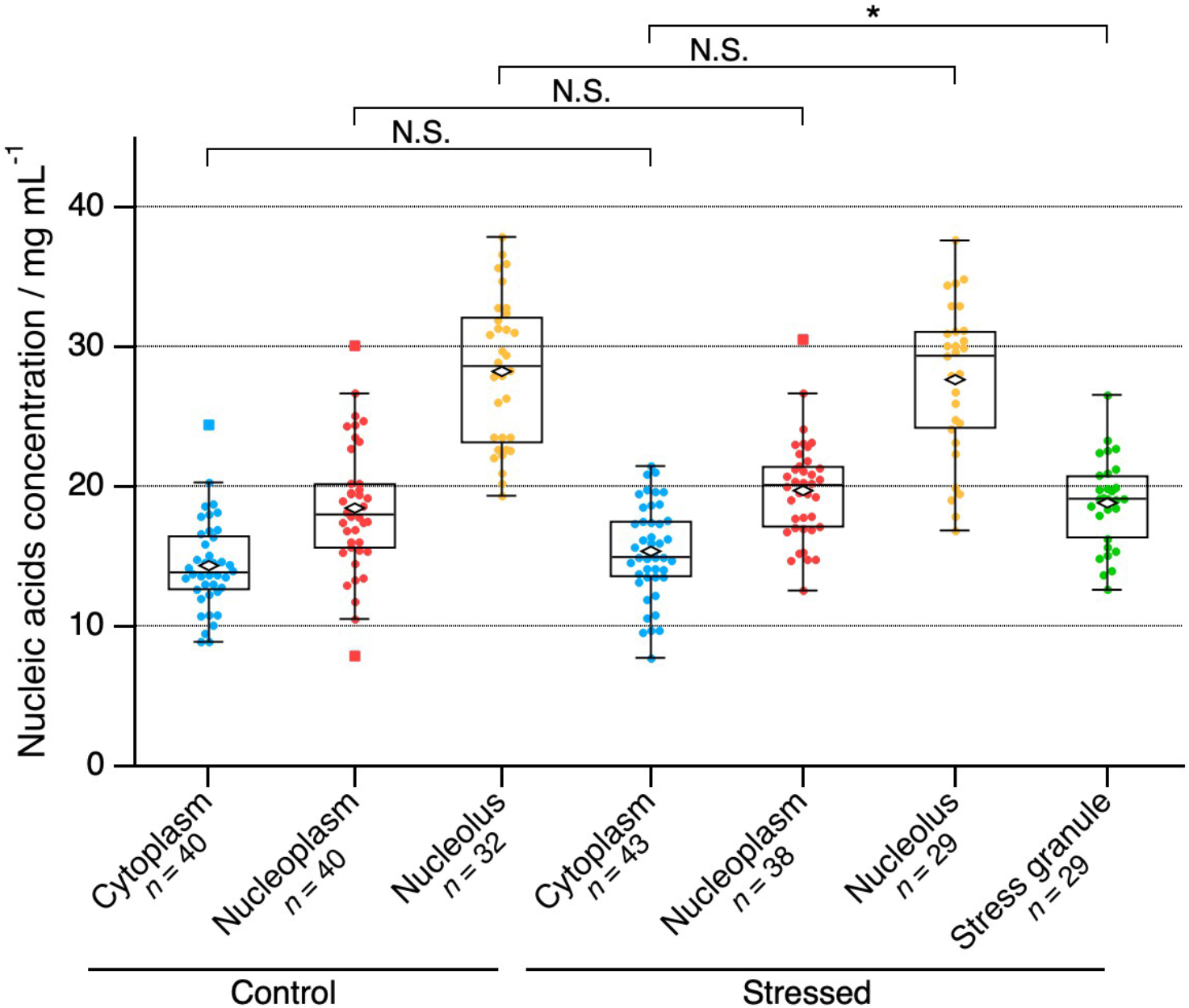
Nucleic acid concentrations of cytoplasm, nucleoplasm, nucleoli, and stress granules in control and stressed cells. Diamond markers are averages and square markers are outliers. *** : *p* < 0.001, N.S.: not significant.

### Liquid Droplets in Intracellular Molecular Crowding Environment

Nucleic acids were confirmed to be concentrated within the SGs; however, this result does not mean that the total concentration of biomolecules in the SGs is higher than in the cytoplasm. The present study indicates that the total biomolecular concentration in the SGs is similar to that in the surrounding intracellular environment exhibiting molecular crowding environment (Figure 2). This result means that an intracellular liquid droplet does not represent a highly-concentrated state, but rather that droplets form via change in the distribution of biomolecules in crowding environment. Thus, the total biomolecular concentration within a droplet is similar to that of the surroundings, and only certain biomolecules are localized. LLPS in a cell is one of the phenomena of intracellular molecular crowding, and the present result exhibits the need to consider a liquid droplet from the viewpoint that LLPS is a localization of specific biomolecules and the resulting droplet is not a dense state with higher biomolecular concentration than the surrounding area.

## Summary

We investigated liquid droplets formed via LLPS in a living cell in terms of the concentration and the local crowding environment. The combination of near-IR fluorescence and Raman imaging offers the visualization of the SGs in living cells and the in situ quantification of the components inside. The Raman spectra of the SGs quantitatively show that the concentration of nucleic acids in the SGs was 20% higher than in surrounding cytoplasmic regions, while the concentration of lipids was lower than surroundings. The net concentration of biomolecules was almost the same in the cytoplasm and the SGs, showing that the crowding environment of the SGs was similar to that of the surrounding cytoplasm. This result means that the SGs are not dense granules formed by very high concentrated biomolecules. The present study shows that intracellular LLPS can be regarded as a phenomenon in which certain biomolecules change their distributions and are localized in molecular crowding environments and the resulting droplets are not a state in which the total biomolecular concentration is higher than that in the surrounding. We also succeeded in tracking cellular responses to oxidative stress in terms of biomolecular concentrations.

Raman imaging allows us to perform the in situ quantitative comparisons of biomolecular concentrations in the SGs and the surrounding cytoplasm in a single living cell. Quantification is essential for an accurate description of the nature and function of intracellular droplets. Raman imaging is a promising method for understanding the nature and function of intracellular LLPS and its relationship to neurodegenerative diseases, leading to the elucidation of the disease mechanisms and the development of their therapy.

## ASSOCIATED CONTENT

### Supporting Information

#### Experimental Procedures

**Figure S1**. Schematic diagram of the experiments.

**Figure S2**. Spectral superimposition of Raman scattering and iRFP fluorescence.

**Figure S3**. Another example of near-IR fluorescence and the corresponding Raman images of stress granules.

**Figure S4**. The intensity of the C–H stretching band relative to that of the O–H stretching band of the cytoplasm and the stress granules

**Figure S5**. The density of water of control and stressed cells.

**Figure S6**. Schematic diagram of the difference between in vitro LLPS and intracellular LLPS.

**Figure S7**. Average Raman spectra of nucleoplasm in control and stressed cells.

**Figure S8**. Average Raman spectra of nucleolus in control and stressed cells.

**Figure S9**. A typical Raman spectrum of an aqueous solution of RNA.

**Figure S10**. A calibration line for RNA concentration.

**Figure S11**. Immunostained fluorescence images of stress granules in HeLa cells.

**Table S1**. Assignments of Raman bands.

## AUTHOR INFORMATION

## Notes

The authors declare no competing financial interests.

## ACKNOWLEDGMENT

This work was supported by JSPS KAKENHI Grant Numbers JP17H05869 (TN), JP20K21468 (TN), JP22H02594 (TN), JP19H05794 (YO), JP19H05795 (YO), JP19H02666 (SK), and JP20H04689 (SK) from the Ministry of Education, Culture, Sports, Science, and Technology in Japan, and JST, PRESTO Grant Number JPMJPR20E5 (SK), CREST and Moonshot Grant Numbers JPMJCR15G2, JPMJCR1852, JPMJCR20E2, JPMJMS2025−14 to YO. SK also acknowledges the Research grant (19E030) from Kurita Water and Environment Foundation (KWEF) in Japan.

## REFERENCES

(1) Hyman, A. A.; Weber, C. A.; Jülicher, F. Liquid-Liquid Phase Separation in Biology. Annu. Rev. Cell Dev. Biol. 2014, 30, 39–58, DOI:10.1146/annurev-cellbio-100913-013325.

(2) Alberti, S.; Gladfelter, A.; Mittag, T. Considerations and Challenges in Studying Liquid-Liquid Phase Separation and Biomolecular Condensates. Cell 2019, 176, 419–434, DOI:10.1016/j.cell.2018.12.035.

(3) Dignon, G. L.; Best, R. B.; Mittal, J. Biomolecular Phase Separation: From Molecular Driving Forces to Macroscopic Properties. Annu. Rev. Phys. Chem. 2020, 71, 53–75, DOI:10.1146/annurev-physchem-071819-113553.

(4) Yoshizawa, T.; Nozawa, R. S.; Jia, T. Z.; Saio, T.; Mori, E. Biological Phase Separation: Cell Biology Meets Biophysics. Biophys. Rev. 2020, 12, 519–539, DOI:10.1007/S12551-020-00680-X.

(5) Wang, B.; Zhang, L.; Dai, T.; Qin, Z.; Lu, H.; Zhang, L.; Zhou, F. Liquid–Liquid Phase Separation in Human Health and Diseases. Signal Transduct. Target. Ther. 2021, 6, 290, DOI:10.1038/s41392-021-00678-1.

(6) Gomes, E.; Shorter, J. The Molecular Language of Membraneless Organelles. J. Biol. Chem. 2019, 294, 7115–7127, DOI:10.1074/jbc.TM118.001192.

(7) Hirose, T.; Ninomiya, K.; Nakagawa, S.; Yamazaki, T. A Guide to Membraneless Organelles and Their Various Roles in Gene Regulation. Nat. Rev. Mol. Cell Biol. 2023, 24, 288–304, DOI:10.1038/s41580-022-00558-8.

(8) Babinchak, W. M.; Surewicz, W. K. Liquid–Liquid Phase Separation and Its Mechanistic Role in Pathological Protein Aggregation. J. Mol. Biol. 2020, 432, 1910–1925, DOI:10.1016/J.JMB.2020.03.004.

(9) Ray, S.; Singh, N.; Kumar, R.; Patel, K.; Pandey, S.; Datta, D.; Mahato, J.; Panigrahi, R.; Navalkar, A.; Mehra, S.; Gadhe, L.; Chatterjee, D.; Sawner, A. S.; Maiti, S.; Bhatia, S.; Gerez, J. A.; Chowdhury, A.; Kumar, A.; Padinhateeri, R.; Riek, R.; Krishnamoorthy, G.; Maji, S. K. α-Synuclein Aggregation Nucleates through Liquid– Liquid Phase Separation. Nat. Chem. 2020, 12, 705–716, DOI:10.1038/s41557-020-0465-9.

(10) Gu, S.; Xu, M.; Chen, L.; Shi, X.; Luo, S.-Z. A Liquid-to-Solid Phase Transition of Cu/Zn Superoxide Dismutase 1 Initiated by Oxidation and Disease Mutation. J. Biol. Chem. 2023, 299, 102857, DOI:10.1016/j.jbc.2022.102857.

(11) Lu, F.-K.; Basu, S.; Igras, V.; Hoang, M. P.; Ji, M.; Fu, D.; Holtom, G. R.; Neel, V. A.; Freudiger, C. W.; Fisher, D. E.; Xie, X. S. Label-Free DNA Imaging in Vivo with Stimulated Raman Scattering Microscopy. Proc. Natl. Acad. Sci. U.S.A. 2015, 112, 11624–11629, DOI:10.1073/pnas.1515121112.

(12) Takahashi, H.; Yanamisawa, A.; Kajimoto, S.; Nakabayashi, T. Observation of the Changes in the Chemical Composition of Lipid Droplets Using Raman Microscopy. Phys. Chem. Chem. Phys. 2020, 22, 21646–21650, DOI:10.1039/D0CP03805A.

(13) Sugimura, T.; Kajimoto, S.; Nakabayashi, T. Label-Free Imaging of Intracellular Temperature by Using the O–H Stretching Raman Band of Water. Angew. Chem. Int. Ed. 2020, 59, 7755–7760, DOI:10.1002/anie.201915846.

(14) Murakami, K.; Kajimoto, S.; Shibata, D.; Kuroi, K.; Fujii, F.; Nakabayashi, T. Observation of Liquid-Liquid Phase Separation of Ataxin-3 and Quantitative Evaluation of Its Concentration in a Single Droplet Using Raman Microscopy. Chem. Sci. 2021, 12, 7411–7418, DOI:10.1039/d0sc06095j.

(15) Yokosawa, K.; Kajimoto, S.; Shibata, D.; Kuroi, K.; Konno, T.; Nakabayashi, T. Concentration Quantification of the Low-Complexity Domain of Fused in Sarcoma inside a Single Droplet and Effects of Solution Parameters. J. Phys. Chem. Lett. 2022, 13, 5692–5697, DOI:10.1021/acs.jpclett.2c00962.

(16) Takeuchi, M.; Kajimoto, S.; Nakabayashi, T. Experimental Evaluation of the Density of Water in a Cell by Raman Microscopy. J. Phys. Chem. Lett. 2017, 8, 5241–5245, DOI:10.1021/acs.jpclett.7b02154.

(17) Shibata, D.; Kajimoto, S.; Nakabayashi, T. Label-Free Tracking of Intracellular Molecular Crowding with Cell-Cycle Progression Using Raman Microscopy. Chem. Phys. Lett. 2021, 779, 138843, DOI:10.1016/J.CPLETT.2021.138843.

(18) Küffner, A. M.; Prodan, M.; Zuccarini, R.; Capasso Palmiero, U.; Faltova, L.; Arosio, P. Acceleration of an Enzymatic Reaction in Liquid Phase Separated Compartments Based on Intrinsically Disordered Protein Domains. ChemSystemsChem 2020, 2, e2000001, DOI:10.1002/syst.202000001.

(19) Avni, A.; Joshi, A.; Walimbe, A.; Pattanashetty, S. G.; Mukhopadhyay, S. Single-Droplet Surface-Enhanced Raman Scattering Decodes the Molecular Determinants of Liquid-Liquid Phase Separation. Nat. Commun. 2022, 13, 4378, DOI:10.1038/s41467-022-32143-0.

(20) Shuster, S. O.; Lee, J. C. Watching Liquid Droplets of TDP-43CTD Age by Raman Spectroscopy. J. Biol. Chem. 2022, 298, 101528, DOI:10.1016/J.JBC.2021.101528.

(21) Choi, S.; Chun, S. Y.; Kwak, K.; Cho, M. Micro-Raman Spectroscopic Analysis of Liquid–Liquid Phase Separation. Phys. Chem. Chem. Phys. 2023, 25, 9051–9060, DOI:10.1039/D2CP05115J.

(22) Kuzmin, A. N.; Pliss, A.; Kachynski, A. V. Biomolecular Component Analysis of Cultured Cell Nucleoli by Raman Microspectrometry. J. Raman Spectrosc. 2013, 44, 198–204, DOI:10.1002/jrs.4173.

(23) Samuel, A. Z.; Sugiyama, K.; Ando, M.; Takeyama, H. Direct Imaging of Intracellular RNA, DNA, and Liquid–Liquid Phase Separated Membraneless Organelles with Raman Microspectroscopy. Commun. Biol. 2022, 5, 1383, DOI:10.1038/s42003-022-04342-4.

(24) Protter, D. S. W.; Parker, R. Principles and Properties of Stress Granules. Trends Cell Biol. 2016, 26, 668–679, DOI:10.1016/j.tcb.2016.05.004.

(25) Hofmann, S.; Kedersha, N.; Anderson, P.; Ivanov, P. Molecular Mechanisms of Stress Granule Assembly and Disassembly. Biochim. Biophys. Acta Mol. Cell Res. 2021, 1868, 118876, DOI:10.1016/J.BBAMCR.2020.118876.

(26) Buchan, J. R.; Parker, R. Eukaryotic Stress Granules: The Ins and Outs of Translation. Mol. Cell 2009, 36, 932–941, DOI:10.1016/J.MOLCEL.2009.11.020.

(27) Bley, N.; Lederer, M.; Pfalz, B.; Reinke, C.; Fuchs, T.; Glaß, M.; Möller, B.; Hüttelmaier, S. Stress Granules Are Dispensable for MRNA Stabilization during Cellular Stress. Nucleic Acids Res. 2015, 43, e26, DOI:10.1093/nar/gku1275.

(28) Wolozin, B. Regulated Protein Aggregation: Stress Granules and Neurodegeneration. Mol. Neurodegener. 2012, 7, 56, DOI:10.1186/1750-1326-7-56.

(29) Anderson, P.; Kedersha, N.; Ivanov, P. Stress Granules, P-Bodies and Cancer. Biochim. Biophys. Acta Gene Regul. Mech. 2015, 1849, 861–870, DOI:10.1016/j.bbagrm.2014.11.009.

(30) Sekiyama, N.; Takaba, K.; Maki-Yonekura, S.; Akagi, K.; Ohtani, Y.; Imamura, K.; Terakawa, T.; Yamashita, K.; Inaoka, D.; Yonekura, K.; Kodama, T. S.; Tochio, H. ALS Mutations in the TIA-1 Prion-like Domain Trigger Highly Condensed Pathogenic Structures. Proc. Natl. Acad. Sci. U.S.A. 2022, 119, e2122523119, DOI:10.1073/pnas.2122523119.

(31) Ellis, R. J. Macromolecular Crowding: An Important but Neglected Aspect of the Intracellular Environment. Curr. Opin. Struct. Biol. 2001, 11, 114–119, DOI:10.1016/S0959-440X(00)00172-X.

(32) Yamamoto, J.; Matsui, A.; Gan, F.; Oura, M.; Ando, R.; Matsuda, T.; Gong, J. P.; Kinjo, M. Quantitative Evaluation of Macromolecular Crowding Environment Based on Translational and Rotational Diffusion Using Polarization Dependent Fluorescence Correlation Spectroscopy. Sci. Rep. 2021, 11, 10594, DOI:10.1038/s41598-021-89987-7.

(33) Schlüßler, R.; Kim, K.; Nötzel, M.; Taubenberger, A.; Abuhattum, S.; Beck, T.; Müller, P.; Maharana, S.; Cojoc, G.; Girardo, S.; Hermann, A.; Alberti, S.; Guck, J. Correlative All-Optical Quantification of Mass Density and Mechanics of Subcellular Compartments with Fluorescence Specificity. eLife 2022, 11, e68490, DOI:10.7554/eLife.68490.

(34) Yang, Q.; Kajimoto, S.; Kobayashi, Y.; Hiramatsu, H.; Nakabayashi, T. Regulation of Cell Volume by Nanosecond Pulsed Electric Fields. J. Phys. Chem. B 2021, 125, 10692–10700, DOI:10.1021/acs.jpcb.1c06058.

(35) Cheung, M. C.; LaCroix, R.; McKenna, B. K.; Liu, L.; Winkelman, J.; Ehrlich, D. J. Intracellular Protein and Nucleic Acid Measured in Eight Cell Types Using Deep-Ultraviolet Mass Mapping. Cytometry A 2013, 83, 540–551, DOI:10.1002/cyto.a.22277.

(36) Theillet, F.-X.; Binolfi, A.; Frembgen-Kesner, T.; Hingorani, K.; Sarkar, M.; Kyne, C.; Li, C.; Crowley, P. B.; Gierasch, L.; Pielak, G. J.; Elcock, A. H.; Gershenson, A.; Selenko, P. Physicochemical Properties of Cells and Their Effects on Intrinsically Disordered Proteins (IDPs). Chem. Rev. 2014, 114, 6661–6714, DOI:10.1021/cr400695p.

