## Supporting Information for "In situ Quantification of Biomolecular Concentration of Cytoplasmic Membraneless Organelles in a Living Cell"

Table of Contents

1. Experimental procedures
2. Figures
3. Table
4. References

### 1. Experimental Procedures

**Cell culture.** HEK293A cells, in which iRFP713-G3BP1 was transfected,[S1] were cultured in a glass-bottom dish (3960-035, 35 mm dish diameter, Matsunami), which was coated with 0.1 mg/L poly-L-lysin, with 2 mL of culture medium (Dulbecco's modified Eagle's medium (DMEM) (D1145, Sigma) supplemented with 10% fetal bovine serum (10437-028, Gibco), 5×104 U/mL penicillin G, 50 mg/L streptomycin sulfate (15070-063, Gibco)), and 200 μg/L hygromycin B (089-0651, Fuji Film). The cells were incubated overnight at 37°C in a 5% CO2 humidified atmosphere. Before the Raman and fluorescence measurements, the culture medium was replaced with Hanks' balanced salt solution (HBSS). To induce oxidative stress and the formation of SGs, cells were further incubated in HBSS containing 0.5 mM sodium arsenite for 30 minutes.[S2] The generation of SGs using this method was confirmed using HeLa cells and an immunofluorescence assay (Figure S11).

**Raman and near-IR fluorescence imaging.** Raman images were obtained with a confocal Raman microscope system (Nanofinder flex2, Tokyo Instruments Inc.) combined with an inverted microscope (Eclipse Ti2, Nikon). The excitation light was 532-nm cw laser beam from a DPSS laser system (Sprout Solo, Lighthouse Photonics). A water-immersion objective lens (Plan Apo IR 60x/NA=1.27, Nikon) was used to excite the sample and collect generated Raman signals. The laser intensity at the entrance of the objective and the exposure time for each image point were 50 mW and 0.1 sec, respectively. Before and after each measurement of a Raman image, fluorescence images were obtained under the epi-illumination condition with a cooled sCMOS camera (CS-53M, Bitran) and an exposure time of 1.5 sec. The excitation and observation wavelengths for the fluorescence measurements were 620 and 720 nm, respectively. All the measurements were performed at room temperature. The IGOR Pro program package (WaveMetrics) was used to analyze Raman and fluorescence images. The singular value decomposition analysis was adopted on each Raman image for noise reduction. Raman images were obtained by mapping the integrated intensity of Raman bands of interest.

**Raman measurements of RNA solution.** Yeast RNA solution (AM7118, Invitrogen) was purchased and used as standard solution for the calibration line. The solution was diluted with ultrapure water. The RNA concentration of the solution was determined by the absorbance at 260 nm. The solution was placed on a glass-bottom dish and Raman spectra of the solution were measured using the Raman microscope.

**Plasmid Availability.** The plasmid to express G3BP1-iRFP713 is deposited to RIKEN DNA bank (RDB17742) and Addgene (#129339).

### 2. Figures

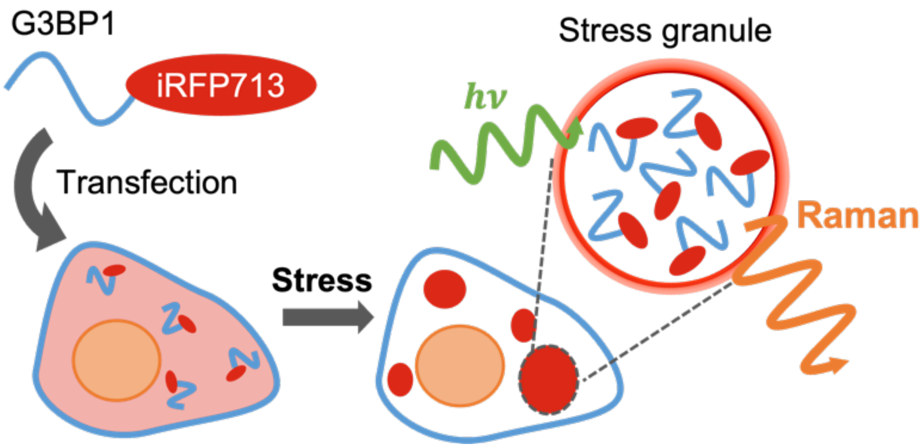


**Figure S1.** Schematic diagram of the experiments. iRFP713 fused G3BP1 protein was expressed in HEK293 cells. Then, cells were exposed to oxidative stress to induce stress granule formation. The positions of the stress granules were determined by near-IR fluorescence images, and Raman images of the stress granules were obtained.

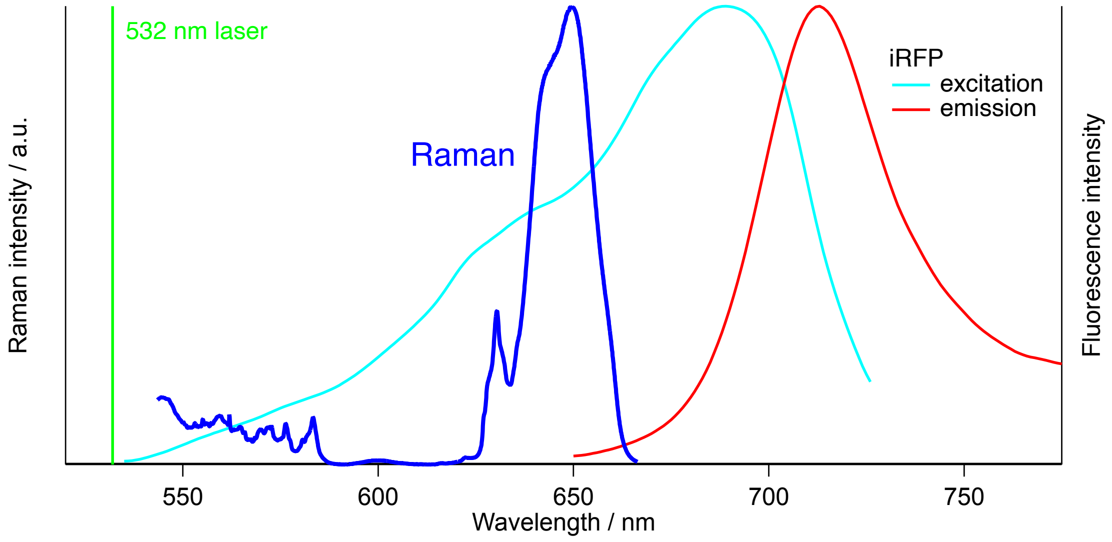


**Figure S2.** Spectral superimposition of Raman scattering obtained by 532 nm excitation light and iRFP fluorescence obtained by visible excitation light. Raman scattering is observed from 540 to 670 nm range. iRFP exhibits fluorescence around 720 nm,[S3,S4] offering a clear window in the visible Raman region.

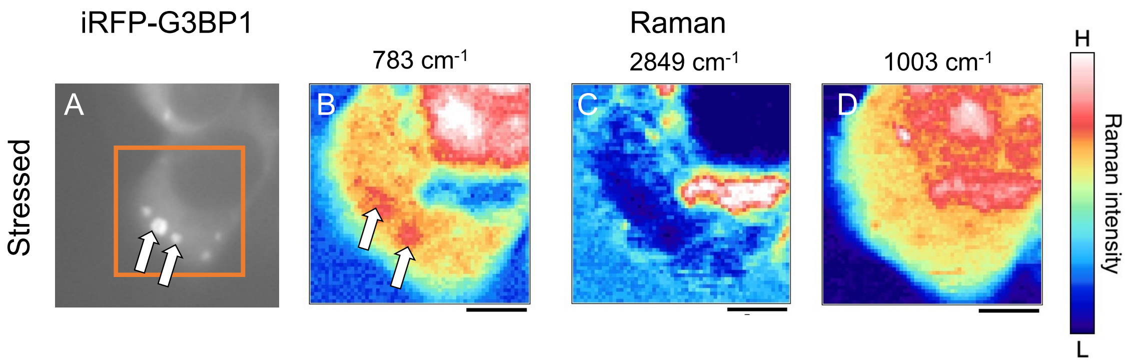


**Figure S3.** Another example of near-IR fluorescence and the corresponding Raman images of stress granules in a cell. An orange box in a fluorescence image shows the region of the Raman imaging. White allows indicate the positions of stress granules.

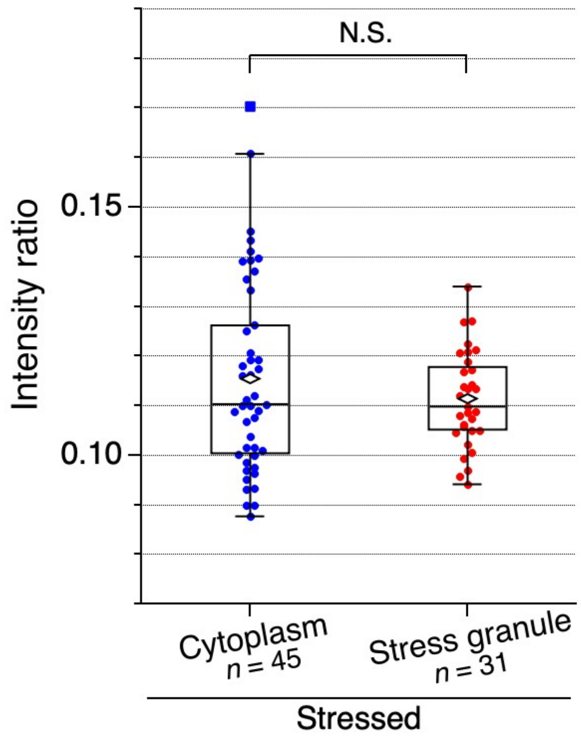


**Figure S4.** The intensity of the C–H stretching band (2800 - 3000 $cm^{-1}$) relative to that of the O–H stretching band (3200 - 3570 $cm^{-1}$) of the cytoplasm and the stress granules in oxidative-stressed cells. Open diamond and closed square marks are the averages and outliers, respectively. N.S.: not significant.

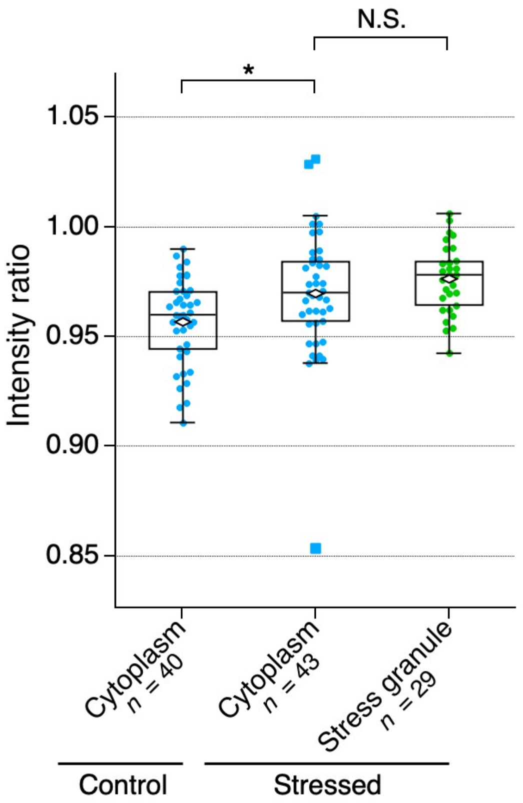


**Figure S5.** The density of water of control and stressed cells. The water density of each compartment was estimated by the intensity ratio between the O–H stretching bands of the intracellular region and the surrounding medium. Diamond markers are averages and square markers are outliers. * : $p < 0.05$, N.S.: not significant. The smaller the intensity ratio, the lower the water density in the local cellular region, i.e., the higher the biomolecular concentration. There was no significant difference in the water densities in the SGs and cytoplasmic regions in stressed cells, showing that they have similar net biomolecular concentrations and crowding environments. Compared with control cells, stressed cells showed higher water density, indicating that the net concentration of biomolecules decreased upon oxidative stress.

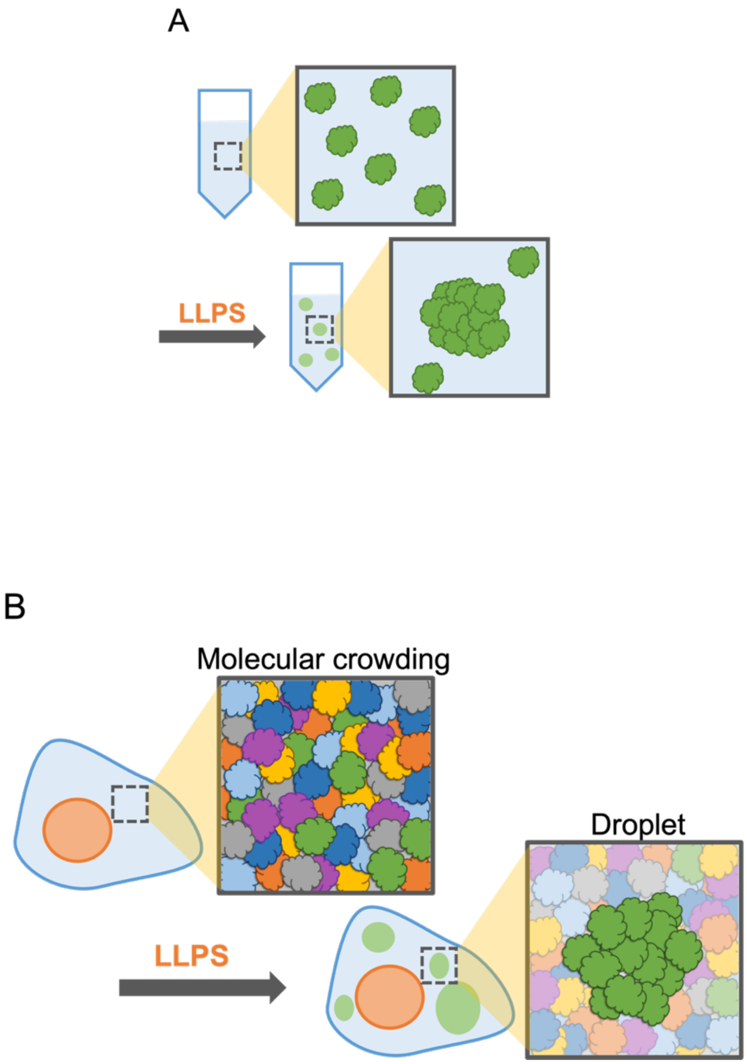


**Figure S6.** Schematic diagram of the difference between in vitro LLPS (A) and intracellular LLPS (B). While in vitro experiments provide a picture of highly concentrated droplets, the concentration of biomolecules in intracellular droplets can be considered to be similar to the surrounding. Only certain biomolecules relating to the formation of droplets, such as G3BP1 and RNA for stress granules, are concentrated, and the others are excluded, keeping the local crowding environments almost constant.

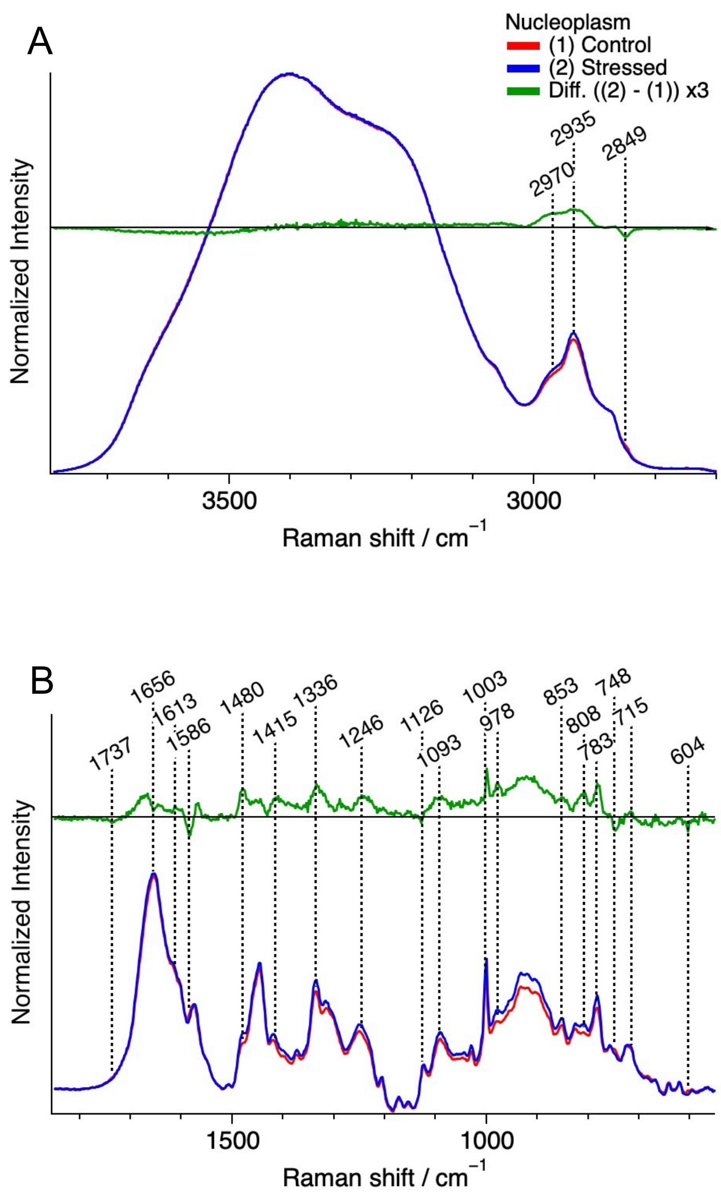


**Figure S7.** Average Raman spectra of nucleoplasm in control (red, $n = 12$) and stressed cells (blue, $n = 13$), and their difference spectrum (stressed – control) (green). Each spectrum was normalized with the integrated intensity of the O–H stretching band (3200 - 3570 $cm^{-1}$). For the fingerprint region, a baseline was obtained by fitting with a fifth-order polynomial and subtracted from each Raman spectrum.

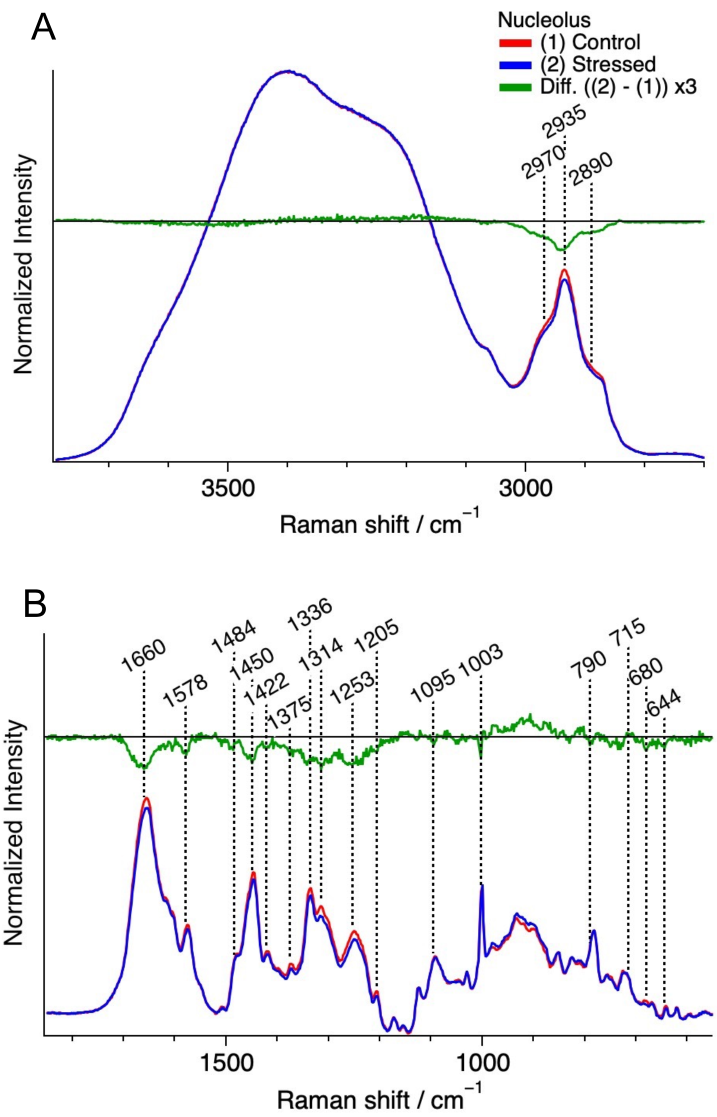


**Figure S8.** Average Raman spectra of nucleolus in control (red, $n$ = 10) and stressed cells (blue, $n$ = 9), and their difference spectrum (stressed ‐ control) (green). Each spectrum was normalized with the integrated intensity of the O–H stretching band (3200 - 3570 $cm^{-1}$). For the fingerprint region, a baseline was obtained by fitting with a fifth-order polynomial and subtracted from each Raman spectrum.

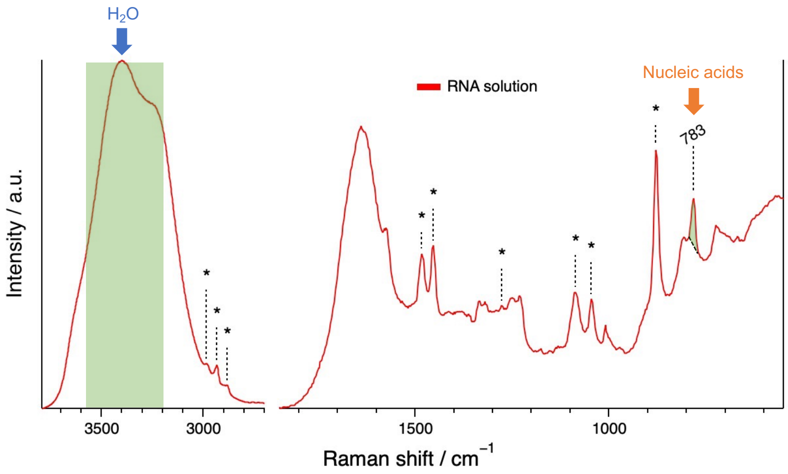


**Figure S9.** A typical Raman spectrum of an aqueous solution of RNA. Asterisks (*) show the Raman bands assigned to ethanol.

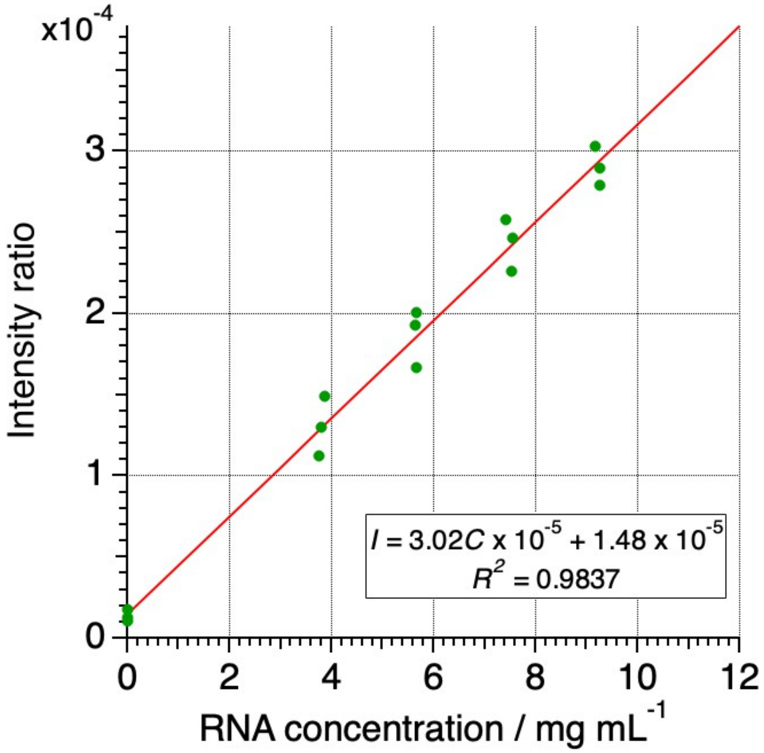


**Figure S10.** A calibration line for RNA concentration obtained by plotting the intensity ratio of the pyrimidine band at 783 $cm^{-1}$ to the O–H stretching band of water at 3200 - 3570 $cm^{-1}$ as a function of RNA concentration.

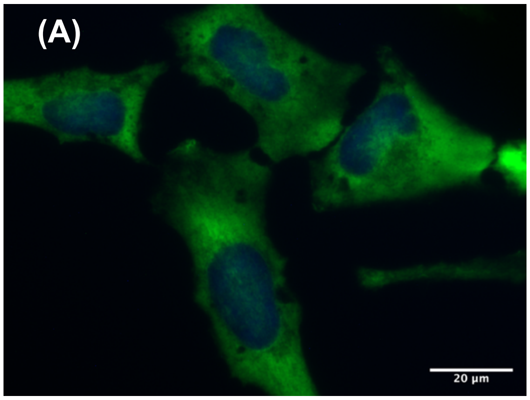


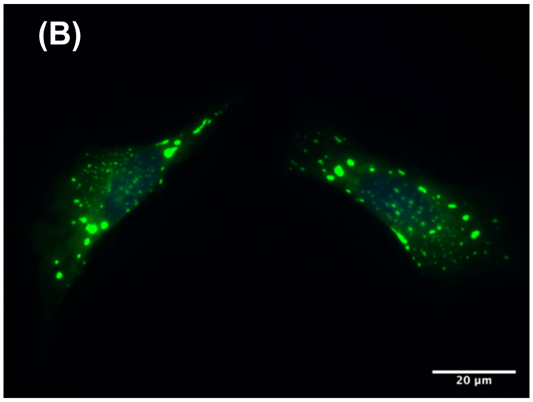


**Figure S11.** Immunostained fluorescence images of stress granules in HeLa cells subjected to oxidative stress: (a) control, (b) stressed cells. Cells were treated with and without 0.5 mM sodium arsenite, and then fixed and immunostained for G3BP1 using G3BP1 polyclonal antibody (Invitrogen, PA5-29455) and donkey anti-rabbit IgG (H+L) highly cross-adsorbed secondary antibody, Alexa Fluor Plus 488 (Invitrogen, A32790). Note that Near-IR fluorescence and Raman images of HEK293 cells shown in the main text were obtained without fixation.

**3. Table**

Table S1. Assignments of Raman bands.[S5, S6, S7]

| Raman shift / $cm^{-1}$ | Assignments | Main molecular components |
|---|---|---|
| 3061 | aromatics | DNAs & RNAs, Proteins |
| 3013 - 3009 | =CH str. | Lipids |
| 2970 | $CH_3$ asym. str. | Proteins, Lipids |
| 2946 - 2935 | $CH_3$ sym. str. | Proteins, Lipids |
| 2890 | $CH_2$ asym. str. | Lipids |
| 2849 | $CH_2$ sym. str. | Lipids |
| 1737 | C=O | Lipids |
| 1660 - 1653 | C=C /Amide I | Lipids / Proteins |
| 1586 - 1584 | | Cytochrome *c* |
| 1578 - 1574 | A, G | DNAs & RNAs |
| 1484 - 1480 | A, G | DNAs & RNAs |
| 1450 - 1445 | CH def. | Proteins, Lipids |
| 1433 | $CH_2$ def. | Lipids |
| 1422 - 1415 | A, G, bk | DNAs & RNAs |
| 1396 | | Cytochrome *c* |
| 1375 | T | DNAs & RNAs |
| 1336 | A, G / CH def. | DNAs & RNAs / Proteins |
| 1314 | G / CH def. | DNAs & RNAs / Proteins / Cytochrome *c* |
| 1302 - 1298 | CH2 tw. | Lipids |
| 1267 - 1262 | Amide III / =CH def. | Proteins / Lipids |
| 1253 | Amide III | Proteins |

| | | |
|---|---|---|
| 1246 | Amide III | Proteins |
| 1205 | Phe, Trp, Tyr | Proteins |
| 1126 | C–N / C–C | Proteins / Lipids/ Cytochrome *c* |
| 1095 - 1093 | $PO_2^-$ | DNAs & RNAs |
| 1003 | Phe. | Proteins |
| 978 - 976 | C–C | Lipids |
| 853 | Tyr | Proteins |
| 808 - 805 | O–P–O | DNAs & RNAs / Lipids |
| 790 - 783 | U, T, C + bk (O–P–O) | DNAs & RNAs |
| 748 | Trp | Proteins / Cytochrome *c* |
| 728 | A | DNAs & RNAs |
| 715 | Choline | Lipids |
| 680 | | Cytochrome *c* |
| 644 | Tyr | Proteins / cytochrome *c* |
| 603 | | Cytochrome *c* |

A; adenine, T; thymine, G; guanine, C; cytosine, U; uracil, Phe; phenylalanine, Tyr; tyrosine, Trp; tryptophan, bk; backbone, def.; deformation, tw.; twist, sym.; symmetric, asym.; asymmetric, str.; stretch.

### 4. References


S1 Yaginuma, H.; Okada, Y. Live Cell Imaging of Metabolic Heterogeneity by Quantitative Fluorescent ATP Indicator Protein, QUEEN-37C. *bioRxiv* **2021**, 2021.10.08.463131, DOI: 10.1101/2021.10.08.463131.

S2 Kedersha, N.; Stoecklin, G.; Ayodele, M.; Yacono, P.; Lykke-Andersen, J.; Fitzler, M. J.; Scheuner, D.; Kaufman, R. J.; Golan, D. E.; Anderson, P. Stress Granules and Processing Bodies Are Dynamically Linked Sites of MRNP Remodeling. *J. Cell Biol.* **2005**, *169*, 871–884, DOI:10.1083/jcb.200502088.

S3 Filonov, G. S.; Piatkevich, K. D.; Ting, L.-M.; Zhang, J.; Kim, K.; Verkhusha, V. V. Bright and Stable Near-Infrared Fluorescent Protein for in Vivo Imaging. *Nat. Biotechnol.* **2011**, *29*, 757–761, DOI:10.1038/nbt.1918.

S4 Lambert, T. J. FPbase: A Community-Editable Fluorescent Protein Database. *Nat. Methods* **2019**, *16*, 277–278, DOI:10.1038/s41592-019-0352-8.

S5 Hobro, A. J.; Rouhi, M.; Blanch, E. W.; Conn, G. L. Raman and Raman Optical Activity (ROA) Analysis of RNA Structural Motifs in Domain I of the EMCV IRES. *Nucleic Acids Res.* **2007**, *35*, 1169–1177, DOI:10.1093/nar/gkm012.

S6 Matthews, Q.; Brolo, A.; Lum, J.; Duan, X.; Jirasek, A. Raman Spectroscopy of Single Human Tumour Cells Exposed to Ionizing Radiation in Vitro. *Phys. Med. Biol.* **2011**, *56*, 19–38, DOI:10.1088/0031-9155/56/1/002.

S7 Hu, S.; Morris, I. K.; Singh, J. P.; Smith, K. M.; Spiro, T. G. Complete Assignment of Cytochrome c Resonance Raman Spectra via Enzymic Reconstitution with Isotopically Labeled Hemes. *J. Am. Chem. Soc.* **1993**, *115*, 12446–12458, DOI:10.1021/ja00079a028.
